# *Legionella pneumophila* translocated translation inhibitors are required for bacterial-induced host cell cycle arrest

**DOI:** 10.1101/479915

**Authors:** Asaf Sol, Erion Lipo, Dennise A. de Jesús, Connor Murphy, Mildred Devereux, Ralph R. Isberg

## Abstract

The cell cycle machinery controls diverse cellular pathways and is tightly regulated. Misregulation of cell division plays a central role in the pathogenesis of many disease processes. Various microbial pathogens interfere with the cell cycle machinery to promote host cell colonization. Although cell cycle modulation is a common theme among pathogens, the role that this interference plays in promoting diseases is unclear. Previously we demonstrated that the G_1_ and G_2_/M phases of the host cell cycle are permissive for *Legionella pneumophila* replication, while S phase provides a toxic environment for bacterial replication. In this study we show that *L. pneumophila* avoids host S phase by blocking host DNA synthesis and preventing cell cycle progression into S phase. Cell cycle arrest upon *Legionella* contact is dependent on the Icm/Dot secretion system. In particular, we found that cell cycle arrest is dependent on the intact enzymatic activity of translocated substrates that inhibits host translation. Moreover, we show that early in infection, the presence of these translation inhibitors is crucial to induce the degradation of the master regulator cyclin D1. Our results demonstrate that the bacterial effectors that inhibit translation are associated with preventing entry of host cells into a phase associated with restriction of *L. pneumophila*. Furthermore, control of cyclin D1 may be a common strategy used by intracellular pathogens to manipulate the host cell cycle and promote bacterial replication.

**Significance:** Recently, we showed that host cell cycle regulatory proteins control *L. pneumophila* growth. In particular, bacterial replication was found to be depressed in S-phase. This indicates that bacterial control of the host cell cycle can limit exposure of the pathogen to antimicrobial events that are cycle-specific. Here we uncovered bacterial factors that induce host cell cycle arrest by inhibiting host protein synthesis and preventing S phase transition. These data are consistent with S-phase toxicity serving as an important antimicrobial response that limits growth of some intracellular pathogens. Moreover, identification of microbial factors that block cell cycle progression and uncovering host cell cycle partners are candidates for future drug development. Our data point to a unifying role of the cell cycle in multiple disease processes.

## Introduction

*Legionella pneumophila* is the causative agent of Legionnaire’s disease (1, 2). The natural hosts of *L. pneumophila* are amoebae, with human disease resulting from aspiration of contaminated aerosols and uptake by alveolar macrophages (1). To sustain intracellular replication, *L. pneumophila* uses the Icm/Dot type IV secretion system (3, 4) which introduces more than 300 Icm/Dot-translocated substrate (IDTS) proteins into the host cell cytosol (5). These IDTS manipulate key host pathways to allow biogenesis of the *Legionella* containing vacuole (LCV; (6-8). Establishment of this replication niche and maintenance of vacuole integrity are two critical steps in supporting intracellular growth and avoiding pathogen recognition by cytosolic innate immune sensing (9). The inability to form the LCV or maintain its integrity result in severe growth defects and rapid pathogen clearance, respectively (10). Our knowledge of host factor restriction of *L. pneumophila* intracellular growth has been greatly enhanced by studies of the targets of the bacterial translocated substrates. For instance, studies on *L. pneumophila* mutants defective for maintaining LCV integrity have allowed significant breakthroughs in identifying the key players in caspase 11-dependent pyroptosis (11).

The eukaryotic cell cycle can be divided into four distinct phases: G_1_, S, G_2_ and M (12). While cells in G_1_ phase commit to proliferation, DNA replication occurs in S phase. Following DNA replication, cells cycle into the G_2_ phase. Transition from G_2_ to M results in new daughter cells. Control of the cell cycle is critical for regulating a number of central processes such as cell differentiation and death, and is tightly controlled by cyclin–dependent Ser/Thr Kinases (CDKs) and their cyclin partners (13). Failure to regulate these proteins in any step of the cell cycle process can lead to catastrophic effects, including uncontrolled cellular growth, such as in cancer (14).

Microbial pathogens can exert cell cycle control on host targets. Notably, a class of proteins called cyclomodulins has been identified that are targeted into the host cell cytosol and interfere with progression through the cell cycle (15, 16). There is also evidence supporting a role for pathogens in modulating tumor progression (17), although the role of such control in supporting disease is still unknown. Recently, studies performed in our laboratory determined that host cell cycle regulatory proteins control *L. pneumophila* growth (18). We demonstrated that the G_1_ and G_2_/M phases of the host cell cycle are permissive for bacterial replication, while S phase provides a toxic environment for bacterial replication. *L. pneumophila* that attempts to initiate replication in S phase shows poor viability, due to a failure to control vacuole integrity that leads to cytosolic exposure of the bacterium and bacterial cell lysis resulting from cytoplasmic innate immune surveillance (11, 18)

Cell cycle progression plays an important role in the intracellular growth of *L.* pneumophil, as emphasized by specific S phase toxicity towards this pathogen (18). Therefore, bacterial control of the host cell cycle can limit exposure of the pathogen to antimicrobial events that are cycle-specific. Indeed, *L. pneumophila* can arrest the host cell cycle which is an effective strategy to avoid S phase toxicity (18, 19). The exact mechanism and the bacterial and host factors that contribute to this cell cycle block remain unknown. Here we show that *Legionella* block of cell cycle progression is dependent on bacterial translocated substrates that interfere with host cell translation. These data provide a mechanism for *L. pneumophila* that allows control of the host cell cycle in multiple cell types.

## Results

### Host cell cycle arrest is dependent on *Legionella* translocated substrates

We previously demonstrated that S phase provides a toxic environment for *Legionella pneumophila* growth and that S phase-infected cells do not progress through the cell cycle after *L. pneumophila* challenge (18). Therefore, avoidance of S has the potential to protect this pathogen from antimicrobial events. To determine if *L. pneumophila* has the capacity to arrest the host cell cycle independently of the phase, we used the double-thymidine block method to synchronize HeLa cells and determine if *L. pneumophila* block cycle progression in a specific phase. Synchronized populations were released from block at time points corresponding to G_1_and G_2_/M and challenged with WT *L. pneumophila-*GFP or a mutant lacking the Icm/Dot type 4 secretion system (T4SS; *dotA3*). Following bacterial challenge, the DNA content of infected cells was compared to that of uninfected cells (Fig. 1A-B, compare GFP^+^, green lines to GFP^-^, black lines) by flow cytometry. The uninfected bystander G_1_- and G_2_/M-synchronized HeLa cells could progress through the cell cycle as measured by accumulation of DNA content over time (Fig. 1A-B, black lines). For instance, bystander G_1_-synchronized cells accumulated DNA by 13 hours post infection (hpi) and by 16 hpi, a G_2_/M population was visible, indicating that these cells progressed through the cell cycle (Fig. 1A, black lines, middle panel). The same pattern applied for G_2_/M synchronized cells that were bystanders at 16 hpi (Figure 1B, black lines, middle panels). In contrast, cells harboring *L. pneumophila* arrested and did not progress through the cell cycle. This was true for both G_1_-and G_2_/M-synchronized cells (Fig. 1A-B, green lines, middle panels). Interestingly, cells infected with the *dotA3* mutant strain progressed normally through the cell cycle. Comparing uninfected cells to those challenged with *dotA3* showed no significant change in DNA content (Fig.1A-B, right panels). Taken together, these data confirm that blocking the host cell cycle is a conserved theme in *L. pneumophila* pathogenesis that relies on the Icm/Dot translocated substrates and is independent of the cell cycle phase encountered by the pathogen.

**Figure 1.**
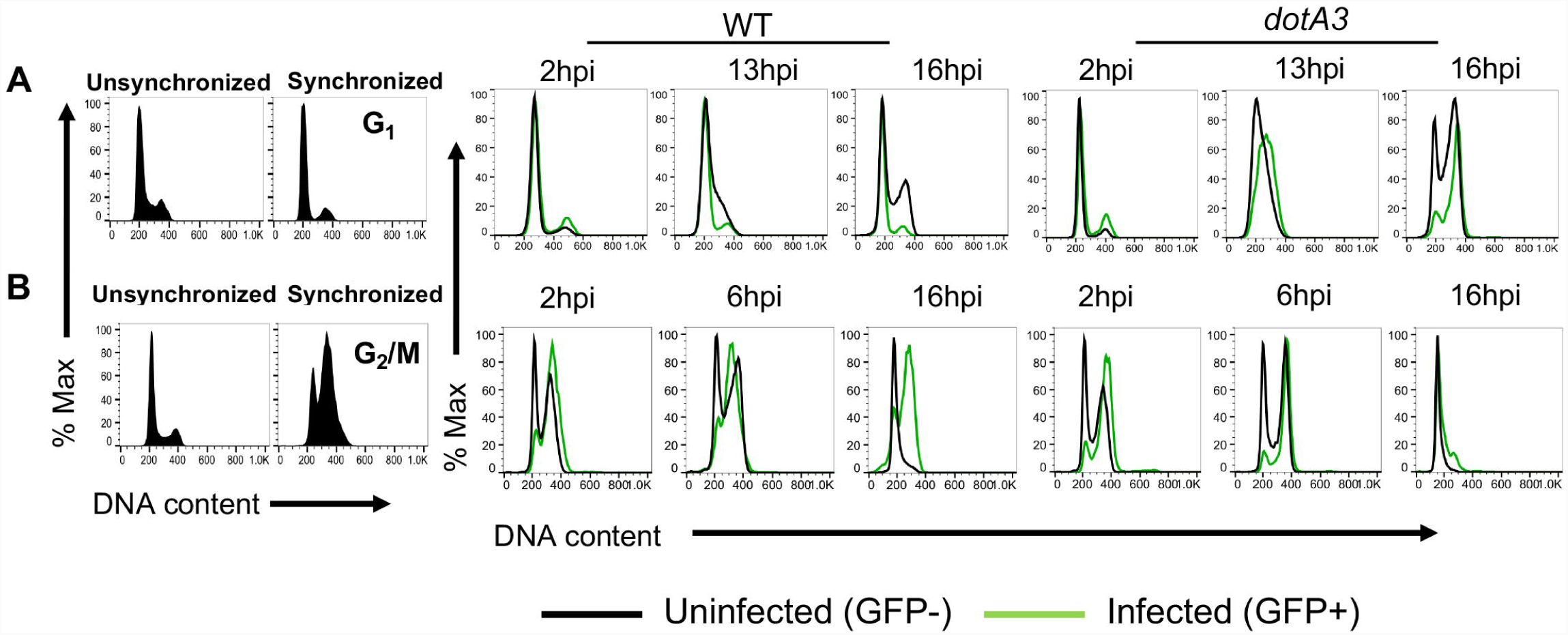
*Legionella pneumophila* Icm/Dot-dependent arrest of the host cell cycle is independent of cell cycle phase. **(A-B)** HeLa cells were synchronized by the double thymidine block method and challenged with WT or *dotA3 L. pneumophila-*GFP. At various times after uptake, cells were collected and analyzed by flow cytometry to determine cell cycle profile (Materials and Methods). (A)11.5hrs and (B) 6.5hrs after release. Left panels with black filled distributions display cell cycle profiles at time of contact with bacteria. Hours post infection: hpi, refers to the total time of contact with bacteria. Uninfected cells (black lines), Infected cells (green lines).

### *Legionella* translocated protein synthesis inhibitors are required for host cell cycle arrest

The Icm/Dot translocated substrates support intracellular growth of *Legionella* in both amoebal species and macrophages and are required for virulence (20-22). Previous studies have shown that protein synthesis in G_1_ phase is a necessary step to progress through the mammalian cell cycle (23) and chemical inhibition of protein synthesis by pharmacological agents such as cycloheximide (CHX) leads to cell cycle arrest (24). *Legionella* has been shown to target host translation elongation through secretion of protein synthesis inhibitors that target host elongation factors eEF1A and eEF1Bγ (25, 26). These inhibitors include the Lgt1-3 glucosyltransferases, SidI and SidL (25, 26). Challenge of host cells with the *L. pneumophila* Δ5 strain, lacking all five of these bacterial protein synthesis inhibitors, results in increased host protein translation compared to cells challenged with WT strain in the first 4 hours of infection (27). To further explore the ability of *Legionella* to block cell cycle progression in a biologically relevant cell type, we determined the proliferation rate of macrophages after *L. pneumophila* challenge by measuring incorporation of the deoxynucleotide analog EdU into newly synthesized DNA. To this end, RAW macrophages were challenged with *L. pneumophila-*GFP strains WT, *dotA3* or Δ*5* and DNA synthesis was measured using EdU incorporation in both the GFP^+^ (infected) and GFP^-^ (bystander) populations (Fig. 2A-B). Macrophages harboring WT *L. pneumophila* showed a large decrease in DNA synthesis compared to cells harboring either the *dotA3* or Δ*5* strains. Strikingly, cells harboring the two mutant strains each had a distinct EdU+ population that overlapped with uninfected cells, indicating a large population of infected cells that show effective cell cycle progression (Fig. 2A-B). Approximately 45% of the uninfected cells showed proliferation, based on EdU incorporation, which was similar to what was observed in response to the *dotA3* mutant. (Fig. 2B). In contrast, approximately 5% of the cell harboring the WT strain showed evidence of proliferation, and this was increased approximately 7 X by introducing the Δ*5* mutation (Fig. 2B).

**Figure 2.**
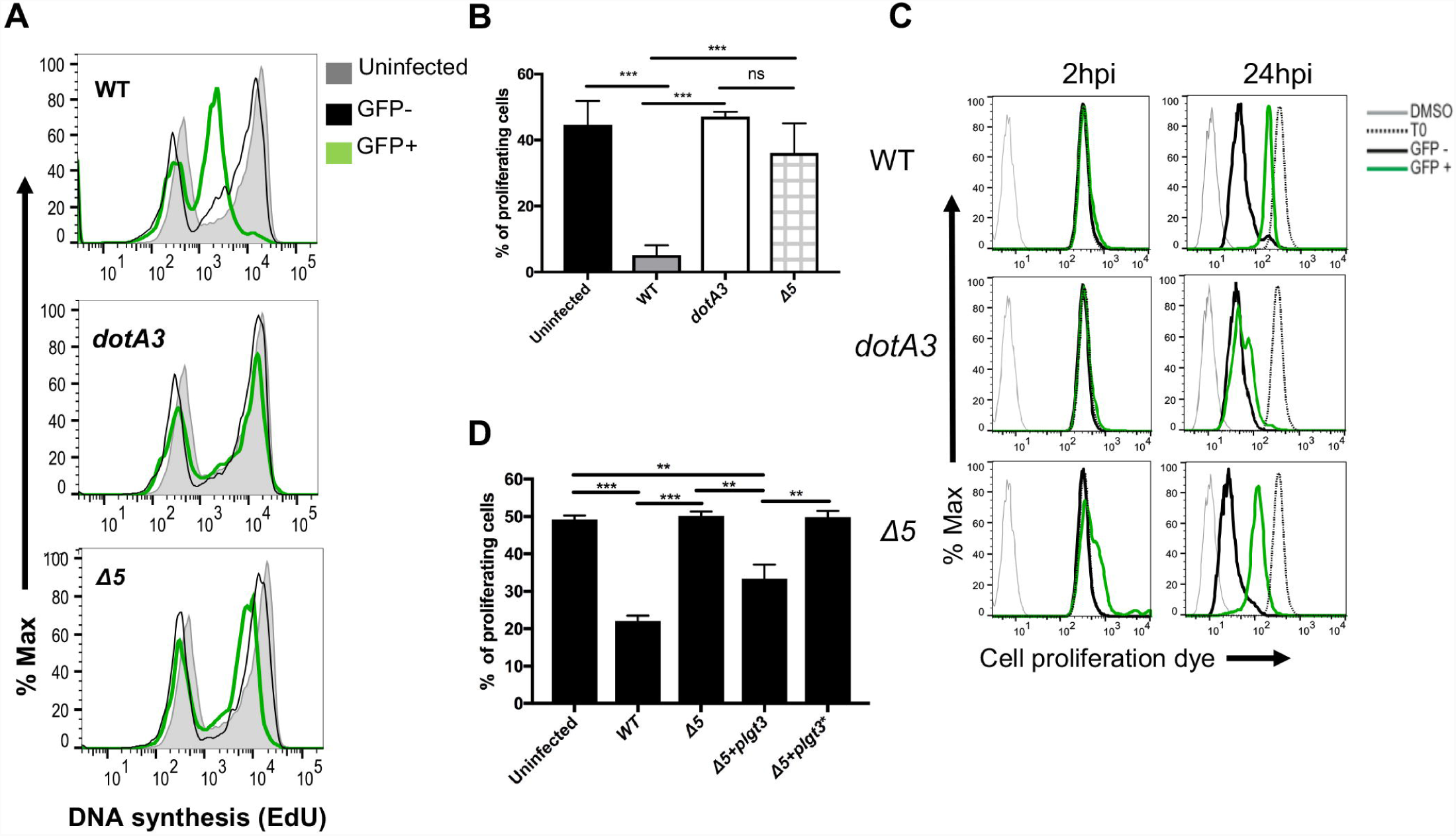
*L. pneumophila* translocated protein synthesis inhibitors are required for the induction of host cell cycle arrest. **(A)** *L. pneumophila* blockade of DNA synthesis requires translocated protein synthesis inhibitors. RAW 264.7 macrophages were challenged with *L. pneumophila* WT, T4SS^-^ or with the Δ5 mutant for 1 hour and then incubated with 20µM EdU for an additional 2 hours. 3 hpi, cells were harvested and analyzed for DNA synthesis using flow cytometry (Materials and Methods). (**B)** Fraction of proliferating cells was quantified using same experimental conditions as A. (**C)** *L. pneumophila* protein synthesis inhibitors block amoebal proliferation. *D. discoideum* cells were incubated with 10µM eFluor^®^ Cell Proliferation Dye. Following staining, cells were challenged with WT, *dotA3* or *L. pneumophila*-GFPΔ5 for 24hrs. At noted time points, the amount of dye within cells was measured by flow cytometry. DMSO: fluorescence of cells with no dye; T0: fluorescence at time of bacterial challenge. (**D)** *L. pneumophila* expressing a single Lgt is sufficient to block host cell proliferation. RAW 264.7 macrophages were challenged with *L. pneumophila* WT or Δ5 complemented with pJB908 vector, Lgt3 or catalytic mutant of Lgt3 (Lgt3*). Following infection, proliferating cells were measured by EdU incorporation and analyzed by flow cytometry. Statistical analyses were performed using unpaired t test. *P < 0.05; **P < 0.01; ***P < 0.001.

The failure of *L. pneumophila* Δ*5* to block cell cycle progression was further verified in the amoebal species, *D. discoideum*, by prelabeling amoebal cells with eFluor cell proliferation dye and measuring dilution of the label as consequence of cell division. Amoebal cells harboring *L. pneumophila* were blocked from proliferation and this block was partially relieved by the Δ*5* mutation (Fig. 2C). The ability to block proliferation of RAW cells dependent on the protein synthesis inhibitors (Fig 2D). The defect in blocking proliferation of RAW macrophages observed with the Δ*5* strain could be reversed by complementation with a plasmid encoding Lgt3 (Fig. 2D). Complementation was not observed in cells challenged with a mutant version of Lgt3 (Lgt3*) that harbored a point mutation in a catalytic residue (Fig. 2D). These data indicate that inhibition of host cell proliferation by *L. pneumophila* requires the presence of translocated protein synthesis inhibitors, and that the catalytic activity of one of these was sufficient to block proliferation.

### *Legionella* effectors trigger the loss of cyclin D

The commitment to start a new round of DNA replication is tightly regulated (28). Transition from G_1_ to S phase is maintained mostly by D-type cyclins and cyclin E, which are controlled by the activity of their partner cyclin dependent kinases (CDKs) (29). These complexes control the activation of a transcriptional network that promotes entry into S phase. D-type cyclins function throughout G_1_ phase while cyclin E shows more activity at the G_1_/S checkpoint. Based on their central role in cell cycle progression, overexpression of D-type cyclins is often associated with cancer, while inhibition leads to cell cycle arrest (30). For this reason, we analyzed the dynamics of cyclin levels in response to *L. pneumophila* challenge of macrophages. The transcription of cyclin D1 (*Ccnd*1) and cyclin E1 (*Ccne1*) genes did not show significant changes when compared to uninfected or *dotA3-*challenged cells (Figure 3A). To confirm that there was no control of these cyclins at the transcriptional level, macrophages were challenged with either the WT or Δ*5* strains, and the response of *Ccnd1* was compared to that of *Egr1*, a gene known to be transcriptionally activated in response to protein synthesis inhibition by *L. pneumophila (*Figure 3B) (31). As a positive control, transcription of *Il6* was analyzed, which is known to respond to *L. pneumophila* independently of the protein synthesis inhibitors (31). While a response to *L. pneumophila* was observed for *Il6* and a protein synthesis inhibitor-dependent response was observed for *Egr1* no significant change was observed in the expression of *Ccnd1* in Δ*5* when compared to WT infected cells (Fig. 3B). Therefore, transcriptional regulation of G_1_ cyclins is an unlikely mechanism for *Legionella* - dependent host cell cycle arrest (Fig. 2).

**Figure 3.**
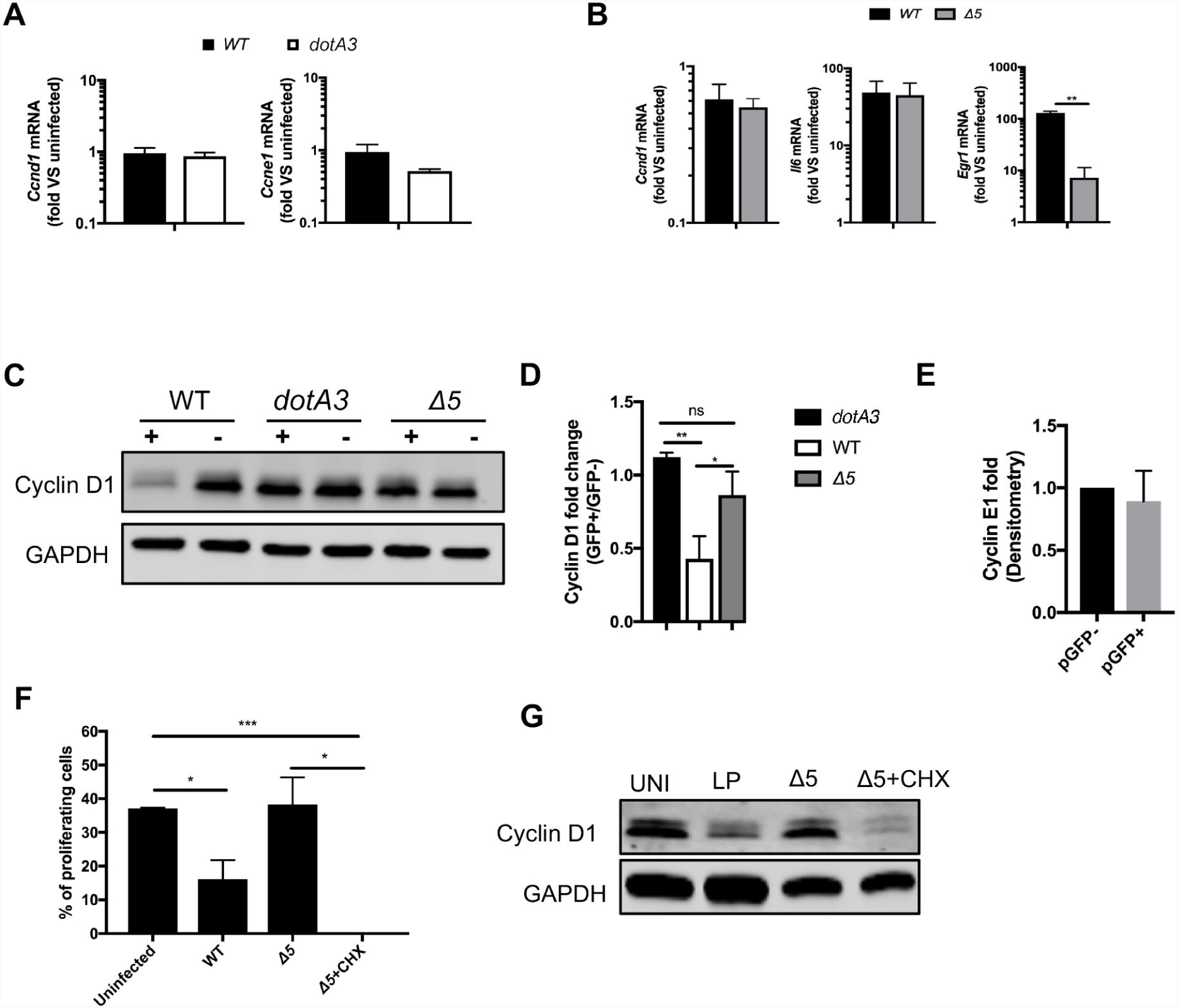
*Legionella* translocated protein synthesis inhibitors induce the rapid degradation of cyclin D1 in infected macrophages. **(A)** Lack of transcriptional effect on cyclin expression. Quantitative PCR analysis of *Ccnd1* and *Ccne1* transcripts in RAW 264.7 macrophages following *L.pneumophila* challenge for 2 hours. Results are normalized based on *Gapdh* expression and presented as relative to uninfected cells. (B) RAW 264.7 macrophages were challenged with *L. pneumophila* WT, *dotA3* or Δ5 for 2 hours and transcript levels of *Ccnd1*, *Il6* and *Egr1* were measured using quantitative PCR analysis. Results presented as relative to uninfected population. (**C-E**) Reduced steady state levels of cyclin D1 in response to *L. pneumophila* requires protein synthesis inhibition. RAW 264.7 macrophages were infected with WT, *dotA3* or Δ5 *L. pneumophila* expressing GFP for 2 hrs and sorted based on fluorescence by flow cytometry. Following infection, the levels of cyclin D1 (C-D) and cyclin E1 (E) in both infected (GFP^+^) and bystanders (GFP^-^) populations were measured by immunoblot. Graph in (D) indicates fold change of cyclin D1 in GFP^+^ compared to GFP^-^ cells. (**F)** RAW 264.7 macrophages were challenged with *L. pneumophila* WT or Δ5 in the absence or presence of 50µg/ml cycloheximide (CHX) for 1 hour and then incubated with 20µM EdU for additional 2 hours. 3 hpi cells were harvested and analyzed for DNA synthesis by flow cytometry. (**G)** RAW 264.7 macrophages were infected with *L. pneumophila* WT or Δ5 as in (F), levels of cyclin D1 in GFP^+^ population were measured by immunoblot. Statistical analyses were performed using unpaired t test. *P < 0.05; **P < 0.01; ***P < 0.001.

We next investigated the role of translational regulation of the G1 cyclins upon *Legionella* infection. Cyclin D1 and cyclin E1 levels were measured in post-sorted cells harboring the *L. pneumophila-*GFP strains (Fig 3C-E, (+)) and compared to bystander cells without bacteria (Fig 3C-E, (-)). Macrophages harboring WT showed loss of cyclin D1 compared to bystander cells (Fig. 3C-D), while cells harboring either *dotA3* or Δ5 showed no appreciable change in cyclin D1 levels (Fig. 3C-D). In contrast, we did not observe significant change in the levels of cyclin E1 protein in cells harboring *Legionella (*Fig. 3E). To further verify that host translation inhibition by *Legionella* is sufficient to induce a proliferation block, we challenged macrophages with Δ*5* in the presence of the translation inhibitor cycloheximide (Fig. 3F-G). Cycloheximide treatment resulted in a complete proliferation block in macrophages challenged with Δ*5 (*Fig. 3F) and was sufficient to recapitulate the loss of cyclin D1 observed in WT *Legionella* infection (Fig. 3G). These results support a model which, cell cycle arrest and destabilization of cyclin D1 by *L. pneumophila* requires the translocation and enzymatic activity of the translation inhibitor effectors.

### A single *L. pneumophila* glucosyltransferase is sufficient to block host cell proliferation

To further study the role of the Lgt proteins in blocking the host cell cycle, we transfected HEK293 cells with plasmids expressing either Lgt3, Lgt1 or catalytically inactive Lgt3 (Lgt3*). To determine effects on translation, the incorporation of the methionine analog azidohomoalanine (AHA) was determined during a two hour labeling, followed by detection via orthogonal linkage to an alkenyl fluorescent probe (32). Cells transfected with either Lgt3 or Lgt1 showed inhibition of protein translation compared to cells transfected with a control vector expressing GFP. In control cells, approximately 43.5% of the cells were in the high translation population compared to 11.1% and 9.59% in the Lgt3 and Lgt1-transfected cells, respectively (Fig. 4A-B). This inhibition was similar to cycloheximide treatment which is a known translation inhibitor (33) and in agreement with previous papers that measured translation rates in cells expressing Lgt proteins using a radiolabeling assay (Fig. 4A-B) (31, 34). Translation inhibition in transfected cells was dependent on Lgt enzymatic activity, as cells transfected with a catalytically inactive Lgt3 (plgt3*) showed translation rates similar to cells harboring a plasmid encoding GFP (42.75% and 43.5% respectively; Fig 4A-B).

**Figure 4.**
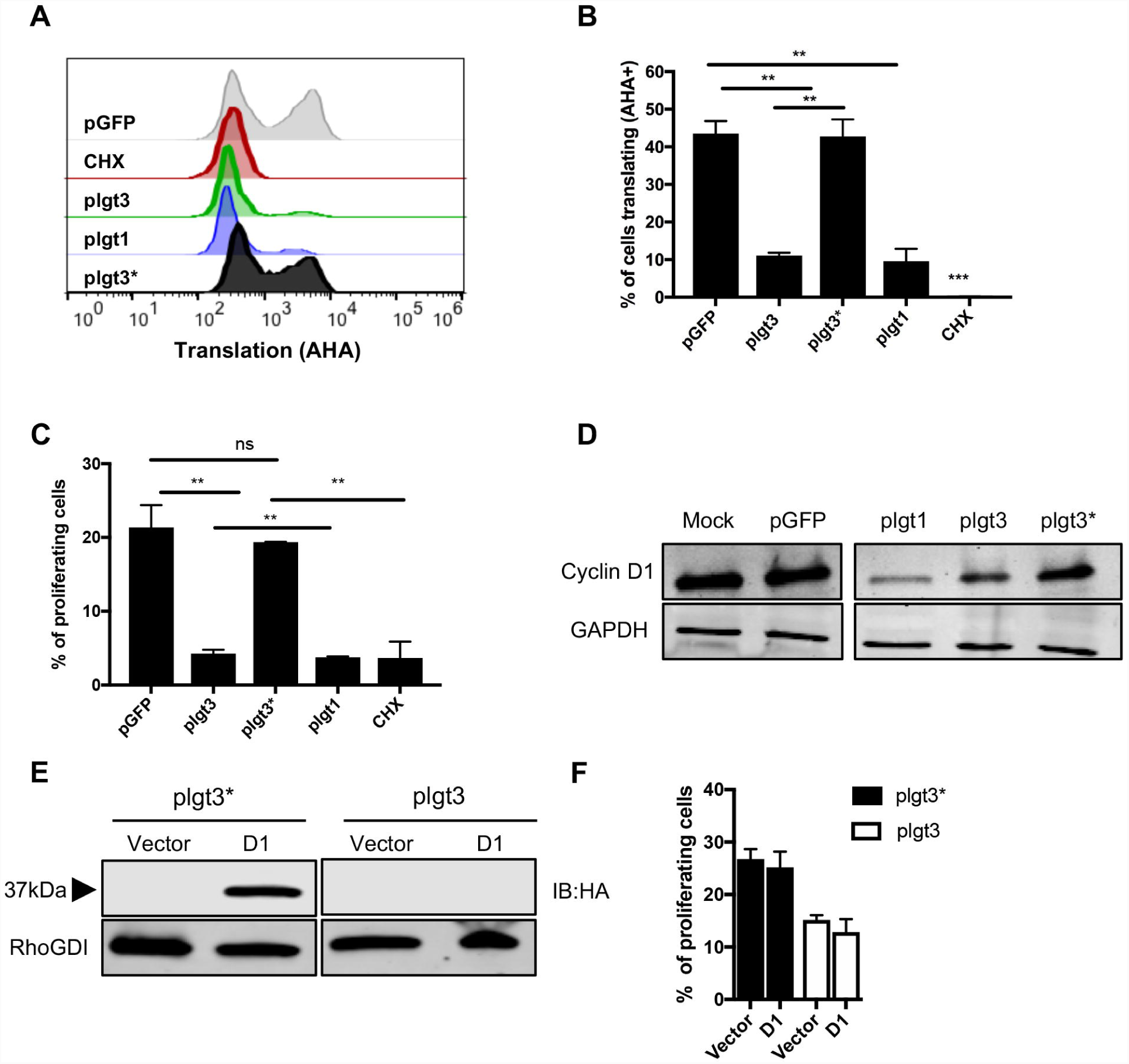
Ectopic expression of the LGTs is sufficient to block host translation and proliferation. **(A-B)** Transfected Lgts block translation. HEK 293T cells were transfected with GFP, Lgt1, Lgt3 or Lgt3* (catalytically dead) for 24 hours. 3 hours prior to harvest, cells were methionine starved for 1 hour and incubated with 50µg/ml cycloheximide (CHX) as appropriate. Cells were then incubated with AHA for 2 hours and *de novo* protein synthesis was measured by orthogonal chemistry and flow cytometry. (**C)** Lgts are sufficient to block proliferation. HEK 293T cells were transfected as in (A). 2 hours prior to harvest cells were incubated with 20µM EdU and *de novo* DNA synthesis was measured as in (B). (**D)** Lowered steady state levels of cyclin D1 in presence of Lgts. HEK 293T cells were transfected as in (A) and cyclin D1 levels were measured by immunoblot. (**E-F)** Overproduced cyclin D1 does not overcome Lgt inhibition. Co-transfection of Lgt3 or Lgt3* inactive enzyme with pcDNA3 harboring HA-cyclin D1 protein. 24 hours post transfection the levels of remaining cyclin D1 were measured by immunoblot (E). (**F)** The percentage of proliferating cells were measured by EdU incorporation as in (C). Statistical analyses were performed using unpaired t test. *P < 0.05; **P < 0.01; ***P < 0.001.

We next tested the link between translation inhibition and cell cycle arrest. To this end, HEK293 was transfected with Lgt1, Lgt3 or the inactive Lgt3* and cell proliferation was measured based on EdU incorporation into DNA and steady state cyclin D1 levels (Fig. 4 C-D). Cells transfected with either Lgt1 or Lgt3 showed a clear cell cycle arrest compared to cells expressing a control GFP or the inactive Lgt3 (Fig 4C). Furthermore, Lgts transfected cells showed decreased expression of cyclin D1 protein (Fig 4D) while cotransfection of HA-cyclin D1 with Lgt3 in HEK293 did not rescue loss of cyclin D1 or the arrest of cell proliferation caused by Lgt3 (Fig 4E-F). These data demonstrated that Lgt3 alone is sufficient to cause cell cycle arrest.

### FZR1 silencing stabilizes cyclin D1 and partially restores DNA synthesis in infected cells

Based on our results showing that overexpression of cyclin D1 did not affect cell cycle arrest in cells expressing *Legionella* translation inhibitors (Fig. 4), we further explored the mechanism of cyclin D1 degradation in cells harboring *L*. *pneumophila*. To determine the rate of cyclin D1 turnover, lysates from infected cells treated with cycloheximide were taken at different time points post-infection and cyclin D1 levels were determined by immunoblotting (Fig. 5A). Cyclin D1 degradation was accelerated in infected cells, resulted in more than 60% reduction in protein levels after 1 hour compared to 20% in uninfected cells (Fig. 5A). Lowered steady state levels were due to proteasomal degradation, based on incubation in the presence of the inhibitor MG132. Treatment with MG132 prior to infection was sufficient to stabilize cyclin D1 in cells harboring WT *Legionella*, as there was no detectable change in cyclin D1 levels in infected cells compared to bystander cells following treatment (Fig. 4B). These results are consistent with the documented role of ubiquitination in regulating cyclin D1 levels (35-37), and in agreement with a previous study demonstrating that MG132 treatment stabilizes cyclin D1 in cells treated with cycloheximide (38).

**Figure 5.**
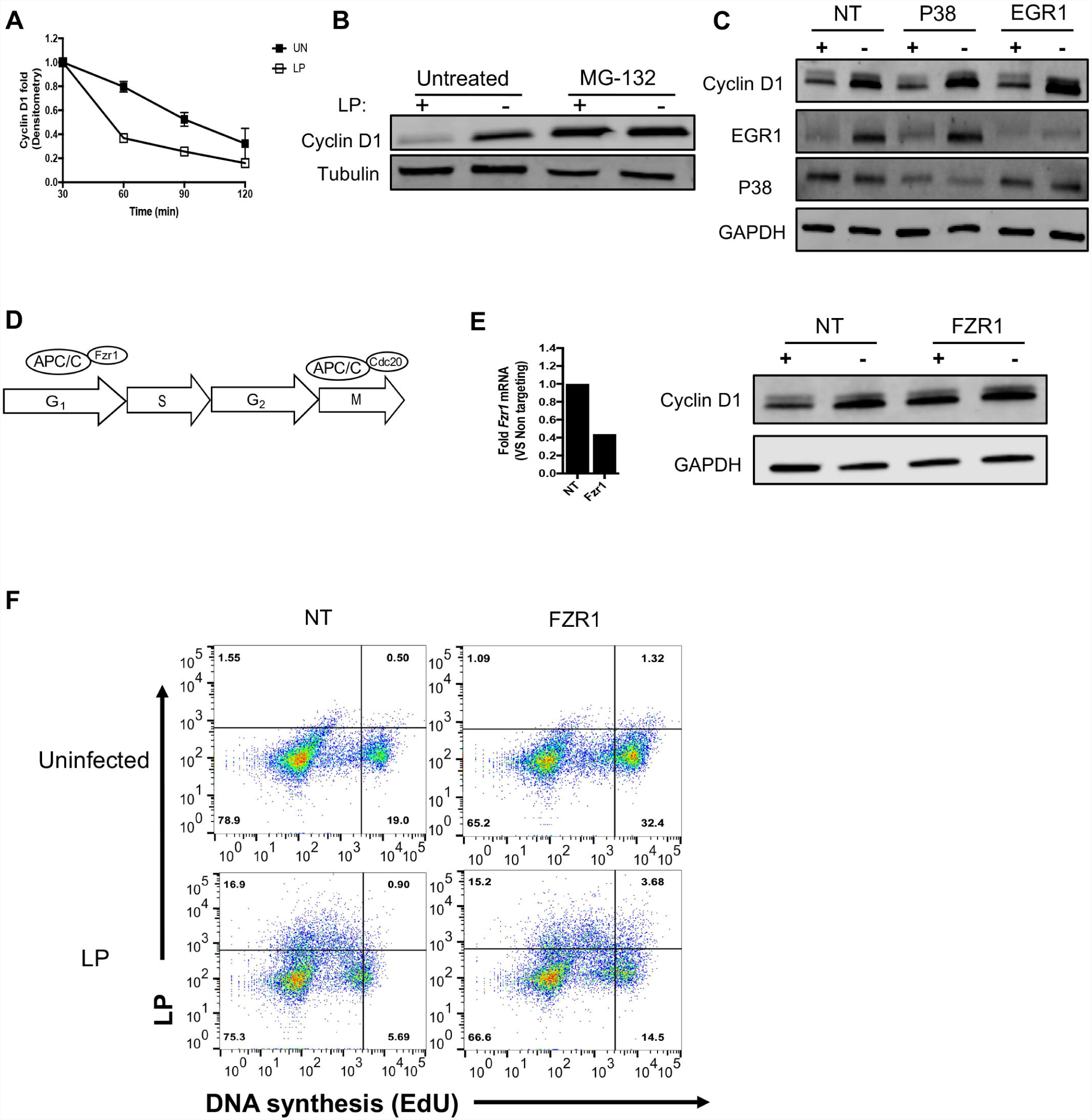
FZR1 silencing partially restores entry into S phase in response to *L. pneumophila*. **(A)** Challenge with *L. pneumophila* destabilizes cyclin D1. Densitometry analysis of cyclin D1 in uninfected RAW 264.7 macrophages and cells challenged with *L. pneumophila* in the presence of 10µg/ml cycloheximide. (**B)** Proteasome inhibition stabilizes cyclin D1. Immunoblot analysis of cyclin D1 in RAW 264.7 macrophages pretreated for 60 min with 10µM of the proteasome inhibitor MG132, challenged with WT *L. pneumophila* for 2 hours and sorted using flow cytometry. (**C)** Immunoblot analysis of cyclin D1 in RAW 264.7 macrophages pretreated with noted siRNA after challenge with *L. pneumophila* WT for 2 hours and sorting. (**D)** Schematic representation of the APC/C association with its coactivators in different phases of the cell cycle. (**E)** FZR1depletion suppresses cyclin D1 destabilization. RAW 264.7 macrophages pre-treated with siRNA against FZR1 or non-targeting siRNA (NT) and analyzed for cyclin D1 as in (C). Insert represent silencing efficiency of targets transcripts. (**F)** Depletion of FZR1partially rescues proliferation of *L. pneumophila-*targeted cells. RAW 264.7 macrophages treated with siRNA against FZR1or Non-targeting siRNA (NT) were challenged with *L. pneumophila* for 1 hour, incubated with 20µM EdU for additional 2 hrs and analyzed for *de novo* DNA synthesis by flow cytometry.

The p38 mitogen-activated protein kinase promotes cyclin D1 degradation and regulates cell proliferation via ubiquitin-dependent turnover of cyclin D1 (39, 40). Furthermore, *Legionella* translation inhibitors activate p38 during infection, providing a potential strategy to drive cyclin D1 degradation (41). For these reasons, we tested if p38 knockdown could restore cyclin D1 levels in infected cells. Surprisingly, p38 silencing did not affect cyclin D1 levels in infected cells (Fig. 5C). In addition, Egr1 silencing, which is another differentially upregulated gene in response to *Legionella* protein inhibitors (Fig. 3B, (31)) showed no effect on cyclin D1 levels in infected cells (Fig. 5C). These data are consistent with a model in which ubiquitination of cyclin D1 in infected cells early in infection is responsible for cyclin D1 turnover.

Levels of cell cycle proteins are maintained through the action of two ubiquitin ligases, the SCF complex (SKP1–CUL1–F-box-protein) and the APC/C (anaphase-promoting complex/cyclosome) complexes, with both targeting specific substrates to the 26S proteasome (42, 43). The APC/C activity is dependent on two co-activators, CDC20 and CDC20-related protein FZR1 (also known as CDH1) (Fig. 5D). While APC^Cdc20^ is important to complete mitosis, association of the APC^FZR1^ complex is important throughout the G_1_, S phase checkpoint (44), with FZR1 knockdowns resulting in early S phase entry and truncated G_1_ phase (45). For this reason, FZR1 was knocked down in macrophages to determine if depletion altered cyclin D1 levels and reduced the cell cycle arrest phenotype after challenge with *L. pneumophila (*Fig. 5E-F). FZR1 knockdown stabilized cyclin D1 protein in infected cells as there was no detectable difference in infected cells compare to bystander cells in FZR1 siRNA treated cells (Fig. 5E). Furthermore, in agreement with previous studies, uninfected cells with FZR1 knockdown resulted in an increase in S phase cells in uninfected population (Fig. 5E, 19.5% to 33.72%). Interestingly, we observed a slight increase in S phase population in infected cells treated with siRNA against FZR1 compared to non-targeting control, consistent with FZR1 playing a role in cell cycle arrest resulting from translation inhibition by *L. pneumophila (*Fig. 5F, lower dot plot). Therefore, we conclude that protein synthesis inhibition by *L. pneumophila* prevents the pool of cyclin D1 from being replenished after it is degraded in an FZR1-dependent fashion.

## Discussion

Manipulation of the host cell cycle is a common strategy used by bacteria and viruses to support intracellular growth (46, 47). Previous work showed that the intracellular pathogen *L. pneumophila* modulates the host cell cycle in a T4S dependent manner (18, 19). In particular, we previously found that *L. pneumophila* growth was enhanced in G_1_ and G_2_, while S-phase was found linked to defective pathogen growth and loss of LCV membrane integrity (18). These results indicate that bacterial control of the host cell cycle can limit exposure of the pathogen to antimicrobial events that are cycle-specific, such as avoiding S phase that is highly restrictive for *L. pneumophila* intracellular growth. It should be noted that the natural hosts of *Legionella*, amoebae, are presumably growth arrested at the time of bacterial contact due to nutrient limitation in their common reservoirs, such as cooling tower. This may provide an opportunity for *L. pneumophila* to lock the host in a state that can promote bacterial growth even when the amoebae respond to nutrient acquisition, such as when they graze on microbial prey. In mammalian hosts, growth arrest is in dynamic conflict with the ability to control intracellular replication, as entry into S phase can be considered an antimicrobial strategy to block intracellular replication. For example, a myeloid precursor responding to growth factor stimulation provides a hostile environment for *L. pneumophila*, as contact with the bacterium during S phase will restrict replication and result in degradation of the LCV with ensuing bacteriolysis.

In this study, we have shown that host cell cycle arrest by *Legionella* is dependent on the Icm/Dot T4SS and is independent of the cell cycle phase at the time of bacterial contact (Fig. 1). In particular, the host proliferation block was found to rely on the activity of five translocated translation inhibitors, with the function of one of these sufficient to cause cell cycle arrest. It should be noted that the *Legionella* glucosyltransferases (Lgts) arrest protein synthesis at the elongation step by glucosylation of eEF1A and eEF1Bγ. We found that early in infection, these translocated substrates are critical for blocks in proliferation and host DNA synthesis (Fig. 2-4). Furthermore, their presence and enzymatic activity induced the rapid degradation of the G_1_/S master regulator, cyclin D1, which prevented cells from entering S phase (Fig. 3-4). Chemical inhibition of protein synthesis by cycloheximide was sufficient to cause cell cycle arrest in the presence of the Δ5 strain lacking the five bacterial protein synthesis inhibitors, indicating that the host translation machinery is likely the *L. pneumophila* effector target that causes arrest (Fig. 3). In addition, ectopic expression of the LGTs was sufficient to block DNA synthesis and led to cell cycle arrest in transfected cells (Fig. 4). These data support a model that the translation block by *Legionella* is the primary cause of host cell cycle arrest, and provides an important selective pressure for retention of this activity. It should be pointed out from previous work, that the cell cycle blockade caused by RNAi depletion of elongation factors confers no cell cycle phase specificity, with the distribution of DNA content in depleted cells appearing similar to proliferating cells (18, 48). This is identical to the result observed in this work. Elongation inhibition clearly provides a strategy to block both entry into S phase as well as prevent mitosis (Fig. 1), which may be similarly restrictive of intracellular growth. As we previously noted, although it seems counterintuitive to encode a protein that allows a block in S phase, in amoeba this phase is extremely short and difficult to identify experimentally (18). The limited opportunities for interaction with the natural host during S phase may also limit the opportunity for negative selection.

There is evidence for a host-induced translation initiation block in response to bacterial infection that is regulated by the mTOR pathway (27, 31, 34, 49). In addition, there may be uncharacterized translocated substrates that interfere with translation initiation. Ribosome profiling analysis of both the wild type *L. pneumophila* strain and a strain lacking all known translocated protein synthesis inhibitors provides solid evidence for other proteins that could interfere with translation initiation (50). Even so, it is clear that the translation blocks provided by these other mechanisms of translation inhibition are not sufficient to interfere with cell cycle progression, based on our analysis of the Δ5 strain (Fig. 1). There is no clear explanation for why an initiation factor block is not sufficient to cause cell cycle arrest. It is known that depletion of translation initiation factors (eIFs) by RNAi treatment causes a dramatic arrest in G1 phase, blocking entry of cells harboring *L. pneumophila* into S phase and stimulating intracellular replication (18, 48). Presumably during *L. pneumophila* growth, the factors that promote translation inhibition allow sufficient breakthrough translation to allow the cell cycle to proceed in their presence.

Control of cyclin D1 function is associated with control of both bacterial and viral growth within hosts, but cyclin activity has different consequences depending on the pathogen (46, 51). A recent study provides evidence that induction of cyclin D1 expression is required for *Salmonella* replication (52), which contrasts strongly with our results that show *Legionella* accelerates cyclin D1 turnover to drive microbial replication (Fig. 5). Cyclin D1 degradation was directly connected to the enzymatic activity of the Lgt proteins (Fig. 3). Due to the tight link between cyclin D1 regulation and the cell cycle state, induction of cyclin D1 degradation by *Legionella* could be the key event that blocks entry into S phase in response to the microorganism. Reversing cyclin D1 destabilization by depleting the FZR1 component of the APC/C complex provided strong circumstantial evidence that the S phase arrest, upon *Legionella* uptake, is a direct result of cyclin D1 destabilization. It was recently shown that depletion of FZR1 could lead to prolonged S phase (45), consistent with the results presented in this communication (Fig. 5). Regulation of cyclin D1 is a key feature in cellular proliferation and as a consequence, cyclin D1 misregulation is found to be involved in several types of cancer (30). Furthermore, manipulation of the APC/C complex is a common strategy used by viruses (54). Regulation of FZR1 activity could provide a rapid strategy to either induce S phase entry or cause cell cycle arrest upon pathogen contact which is a common theme in many pathogenic events.

In summary, our results demonstrate that specific *Legionella* translocated substrates interfere with the host cell cycle causing cells to growth arrest at the stage in the cell cycle that encounters the microorganism. These data provide evidence for the key role for cyclin D1 turnover in infected cells and argue that the APC/C complex promotes bacterial replication by causing cyclin D1 degradation and preventing G_1_ cells from entering S phase, which restrict *L. pneumophila* growth. Although *Legionella* infects terminally differentiated human macrophages, recent studies show that tissue resident macrophage can proliferate *in situ* in response to different triggers, providing potentially antimicrobial reservoirs (55, 56). These data point to a model in which S-phase re-entry by macrophages harboring *Legionella* could serve as a poorly appreciated strategy to promote restriction of selected pathogens. Future work will be devoted to study the interplay between the cell cycle machinery and *Legionella* during disease, and in particular, the role of the APC/C complex in modulating these interactions.

## Materials and Methods

### Bacteria and culturing media

*L. pneuomophila* Philadelphia 1 strains used in this study are described in Table 1. *L. pneuomophila* was grown in ACES-buffered yeast extract (AYE) broth medium supplemented with 100 µg/ml thymidine and maintained on solid medium using buffered charcoal yeast extract agar (BCYE). Strains carrying pGFP Cm^R^ plasmids were maintained on BCYE agar containing 100 µg/ml thymidine and 5 µg/ml chloramphenicol. When appropriate, IPTG was added to a final concentration = 1 mM. Strains carrying pJB908 plasmid were grown in AYE without thymidine supplementation.

**Table 1.** Plasmids and bacterial strains used in this study.

| Plasmid/Strain | Genotype/Characteristics | Reference |
| --- | --- | --- |
| **pGFP** | oriRSF101Cm<sup>R</sup>ptac::GFP<sup>+</sup> | (27) |
| **Lp01** | Philadelphia 1, wild-type strain | (18) |
| ***dotA3*** | T4SS translocation deficient | (18) |
| **Δ*flaA*** | Lp02 Δ*flaA* | (27) |
| ***dotA3* Δ*flaA*** | Type IV secretion system deficient, Δ*flaA* | (27) |
| **Δ5 Δ*flaA*** | Missing 5 translocated effectors required for translation inhibition: Lgt1-3, SidI and SidL, Δ*flaA* | (27) |
| **Δ*flaA*-pGFP** | Δ*flaA* containing pGFP | (27) |
| ***dotA3* Δ*flaA*-pGFP** | *dotA3* Δ*flaA* containing pGFP | (27) |
| **Δ5 Δ*flaA*-pGFP** | Δ5 Δ*flaA* containing pGFP | (27) |
| **pJB908** | ptac *thyA*<sup>+</sup>, IPTG inducible | (59) |
| **Δ5 Δ*flaA*-pJB908** | Δ5 Δ*flaA* containing pJB908 | This study |
| **Δ5 Δ*flaA*-plgt3** | Δ5 Δ*flaA*-plgt3, Lgt3 complementation | This study |
| **Δ5 Δ*flaA*-plgt3\*** | Δ5 Δ*flaA*-plgt3, Lgt3 complementation with catalytic dead Lgt3 | This study |
| **pT-REx-DEST31** | Gateway vector, Mammalian expression, N-terminal His tag | This study |
| **plgt3** | Lgt3 expression vector, pT-Rex-DEST31 backbone | This study |
| **plgt3\*** | Catalytic dead Lgt3 expression vector, pT-Rex-DEST31 backbone | This study |
| **plgt1** | Lgt1 expression vector, pT-Rex-DEST31 backbone | This study |
| **pD1-HA** | pcDNA cyclin D1-HA | (58) |

### Mammalian cell culture

RAW 264.7 cells (ATCC) were grown in RPMI supplemented with 10% heat-inactivated FBS and 1 mM L-glutamine. HeLa and HEK293T cells were cultivated in Dulbecco modified Eagle medium ((DMEM; Gibco) supplemented with 10% heat-inactivated FBS (Gibco) All cell lines were cultured at 37°C with 5% CO2.

### Infection with *Legionella pneumophila*

*L. pneumophila* strains were grown on patches on CYE plates36-48h prior to infection. Patches were then resuspended in AYE media with the appropriate supplements and grown overnight. Challenge of host cells was carried out at a multiplicity of infection (MOI) = 50 to allow for pool adhesion by the Δ*flaA* strains. Following challenge, plates were subjected to centrifugation for 10 min at 400 × g to synchronize the infection. 1-2 hours after uptake, cells were washed twice to remove extracellular bacteria, and resuspended in fresh medium.

### HeLa cell cycle synchronization

HeLa cells were synchronized by the double thymidine block (57). Briefly, 1 × 10^6^ HeLa cells were incubated with an excess of 2 mM thymidine for 18h. Cells were washed two times with 1 X PBS and released by incubation for 8 hours in DMEM-FBS without thymidine. Following release, cells were treated with a second dose of 2 mM thymidine. After 14 to 16 h, cells were collected and re-plated at 2.5 × 10^5^ cells/well in 6-well plates for either cytometry or for challenge with *L. pneumophila* at the indicated times after release.

### EdU incorporation and *de novo* DNA synthesis detection

RAW 264.7 cells were plated at 1 × 10^6^ cells/well in 6-well plates and then challenged with *L. pneumophila* at MOI = 50. 1hr post infection (hpi), cells were washed with 1X PBS and incubated with medium containing 20µM of the deoxynucleotide analog EdU for an additional 2 hours. 3hpi, cells were collected and *de novo* DNA synthesis was measured by orthogonal Click-iT chemistry using Click-iT™ EdU Alexa Fluor™ 647 Assay Kit (Invitrogen) and analyzed by flow cytometry using BD LSRII analyzer.

### Proliferation assay in *D. discoideum*

To measure cell proliferation in amoebae at 23C°, stationary phase *D. discoideum* AX4 cells were collected and seeded at 2 × 10^6^ cells/well on 6-well plates in PBS. Cells were incubated in the dark with 10µM Cell Proliferation Dye (eBioscience) or DMSO equivalent for 30 minutes. Uninternalized dye was washed away with ice cold 1% BSA in 1X-PBS and replaced by HL-5 medium (18). Cells were collected at the indicated times and analyzed by flow cytometry using BD LSRII analyzer.

### Immunoblotting

RAW 264.7 cells were plated at a density of 1 × 10^7^ cells/well in 10 cm dishes and then challenged with *L. pneumophila*-GFP strains at MOI = 50. When appropriate, cells were incubated for 1 hour prior to *Legionella* challenge with 50µg/ml cycloheximide (CHX) or 10µM MG132. 2 hpi, cells were washed with 1X PBS and harvested. Cells were fluorescently sorted based on GFP signal using BD INFLUX sorter. Post-sorted infected cells (GFP^+^) and bystander cells (GFP^-^) were centrifuged and lysed using 2 X SDS Laemmli sample buffer (0.125 M Tris-HCl (pH = 6.8), 4% SDS, 20% glycerol, 10% 20% glycerol, 10% β-mercaptoethanol, 0.01% bromophenol blue) and boiled for 10 min. Proteins were transferred to PVDF membranes, blocked in 5% BSA and probed with antibodies to cyclin D1 (Abcam), cyclin E1 (Cell Signaling), P38 (Cell Signaling), EGR1 (Cell Signaling), RhoGDI (Cell Signaling), GAPDH (Cell Signaling) or Tubulin (Sigma). Immunoblotting with primary antibodies was carried overnight in 4°. Dylight anti-rabbit IgG 680 and anti-mouse IgG 800 were used as secondary antibodies (Cell Signaling, 1:20000). Capture and analysis of membranes was performed using the Odyssey scanner and the Image Studio software (LI-COR Biosciences).

### Detection of translation and quantification

HEK-293T cells were seeded at 2 × 10^6^ cells per well in a 6-well plate. Cells were transfected using Lipofectamine 2000 (Invitrogen) according to the manufacturer’s protocol. For detection of whole cell translation, after 24 hours post transfection, the medium was changed to Methionine Free Media for one hour. Cells were then labeled with 100µM of L-azidohomoalanine (AHA, Invitrogen). After 2 hours cells were washed with PBS and fixed with 4% paraformaldehyde for 15mins. Cells were then incubated with 50µM of Dylight650 conjugated phosphine reagent (Pierce) at 37°C for 3 hours. Cells were extensively washed to remove excess dye with 0.5% Tween-20/PBS and analyzed by BD LSR II flow cytometer.

### Quantitative RT-PCR

RAW 264.7 cells were plated at 1 × 10^7^ cells/well in 10 cm dishes and then challenged with *L. pneumophila-*GFP at MOI = 50. 3 hpi, cells were washed with 1X PBS and harvested. Cells were based on GFP signal using BD INFLUX sorter. Post-sorted infected cells (GFP^+^) and bystander cells (GFP^-^) were subjected to centrifugation and resuspended in RLT buffer according to manufacturer’s instruction (Qiagen). RNA was isolated using the RNAeasy kit (Qiagen) and cDNA synthesis was performed using SuperScrip IV VILO Master Mix (Invitrogen). Transcripts were measured using SYBR Green (Applied Biosystems) from cDNA templates using the following primer pairs: *Ccnd1(*5’ TGCCGAGAAGTTGTGCATCTA and 3’ ACCTTGACGAAGACCACTTGT). *Ccne1 (*5’ CTGTGAAAAGCGAGGATAGCA and 3’ AAAGTAGGGGTGGGGATTGT). *Il6 (*5’ CGATGATGCACTTGCAGAAA and 3’ AAGGAGACCAGAAGACCTCA). *Egr1 (*5’ ACAACCCTATGAGCACCTGAC and 3’ CCTCTGCTCAATAGGGTCGG). *Gapdh (*5’ CAAGGTCATCCCAGAGCTGAA and 3’ ACACAGGCAGCACCTAGAC). Gene expression levels were quantified using the 2^-ΔΔCt^ method with *Gapdh* as endogenous control. All reactions were carried out using the StepOnePlus Real-Time PCR system (Applied Biosystems).

### Gene Silencing by Small Interfering RNA (siRNA)

All siRNA transfection experiments were performed using lipofectamine RNAiMax according to manufacturer’s instructions for 72h (Life Technologies). siRNA oligonucleotides were purchased from Dharmacon and were specifically designed to deplete mouse FZR1, P38 and Egr1. Non-targeting siRNA was used as a control.

### Transfection of *Legionella* glucosyltransferase

HEK 293T cells were seeded at 2 × 10^6^ cells per well in a 6-well tissue culture plate. Cells were transfected using Lipofectamine 2000 or 3000 (Invitrogen) according to the manufacturer’s protocol. At 24 hpi, cells were lysed in Laemmli Sample Buffer (Biorad) for Western blot analysis or labeled for translation or DNA synthesis using Click-iT chemistry as described above. In co-transfection assays we used pcDNA Ccnd1 with a C terminal HA tag. Plasmid was a gift from Dr. Bruce Zetter ((58), Addgene plasmid # 11181).

## Acknowledgments

We thank Stephen Kwok and Allen Parmelee from Tufts University flow cytometry core for their technical assistance in sorting experiments. This work was supported by HHMI and NIAID grant R01AI113211 to R.R.I. AS was supported by HHMI. DADJ was supported by NIH Post-Baccalaureate Research Education Program Grant 5R25GM066567, NIH Training Grant T32AI07422 and a Predoctoral Fellowship 1F31AI098423-01 from the National Institute of Allergies and Infectious Diseases. EL and DADJ were supported by NIH training grant T32GM07310.

